# Surprise-induced enhancements in the associability of Pavlovian cues facilitate learning across behavior systems

**DOI:** 10.1101/2021.06.14.448382

**Authors:** Inmaculada Márquez, Gabriel Loewinger, Juan Pedro Vargas, Juan Carlos López, Estrella Díaz, Guillem R. Esber

## Abstract

Surprising violations of outcome expectancies have long been known to enhance the *associability* of Pavlovian cues; that is, the rate at which the cue enters into further associations. The adaptive value of such enhancements resides in promoting new learning in the face of uncertainty. However, it is unclear whether associability enhancements reflect increased associative plasticity within a particular behavior system, or whether they can facilitate learning between a cue and any arbitrary outcome, as suggested by attentional models of conditioning. Here, we show evidence consistent with the latter hypothesis. Violating the outcome expectancies generated by a cue in an appetitive setting (feeding behavior system) facilitated subsequent learning about the cue in an aversive setting (defense behavior system). In addition to shedding light on the nature of associability enhancements, our findings offer the neuroscientist a behavioral tool to dissociate their neural substrates from those of other, behavior system- or valence-specific changes. Moreover, our results present an opportunity to utilize associability enhancements to the advantage of counterconditioning procedures in therapeutic contexts.

---

In an ever-changing world, brain mechanisms have evolved to modulate the associability of Pavlovian cues in order to meet the learning demands of the environment (e.g., Mitchell & Le Pelley, 2010). One form of modulation is captured by the so-called *uncertainty principle*, according to which a cue’s associability increases whenever its consequences are surprising (Pearce & Hall, 1980; Pearce et al., 1982). In support of this notion, cues that predict an outcome inconsistently (i.e., partial reinforcement) are subsequently learned about more rapidly than cues that predict the outcome consistently (i.e., continuous reinforcement; Haselgrove et al., 2010, Collins & Pearce, 1985). Similarly, repeated confirmation of outcome expectancies decreases a cue’s associability (Pearce & Hall, 1979; Griffiths et al., 2011; Mackintosh & Turner, 1971), whereas a sudden violation of those expectancies restores it (Hall & Pearce, 1982; Dickinson et al., 1976; Holland, 1984). Surprise-induced associability enhancements have been documented both in appetitive (e.g., Holland, 1984) and aversive (e.g., Dickinson et al., 1976) procedures as well as across phylogenetically distant species (e.g., rats: Kaye & Pearce, 1984; pigeons: Collins & Pearce, 1985; humans: Hogarth et al., 2008; Russo et al., 2019), suggesting they might constitute a widespread, if not universal property of learning systems.

While these findings have fostered important neurobiological discoveries (reviewed in: Holland & Schiffino, 2016; Roesch et al., 2012; Holland & Maddux, 2010) and theoretical developments (Dayan et al., 2000; Le Pelley, 2004; Courville et al., 2006; Pearce & Mackintosh, 2010; Esber & Haselgrove, 2011), the nature and scope of associability enhancements remains poorly understood. On the one hand, such enhancements might reflect a state of heightened associative plasticity involving a specific association or behavior system (e.g., feeding, mating, defense, etc.; Timberlake, 1993; 1994; Cabrera et al., 2019). Such a labile state would facilitate the updating of associative representations involving the cue and outcomes within that behavior system. It follows from this view that a surprise-induced associability enhancement by a food-predictive cue, for instance, should translate as more rapid learning between that cue and food-related outcomes (including food omission), but not necessarily outcomes related to other behavior systems, such as the presence of a sexual partner (Domjan & Gutiérrez, 2019). On the other hand, associability enhancements might arise from increased attentional processing of the cue, as assumed by attentional models of associative learning (Mackintosh, 1975; Pearce & Hall, 1980). If so, those enhancements should manifest as faster learning regardless of the nature of the outcome and the behavior system engaged. Since studies on associability modulation have traditionally employed a single reinforcer or reinforcer class (thus engaging a single behavior system), this issue remains unresolved.

To decide between these alternatives, we violated the outcome expectancies generated by a cue in an appetitive setting (feeding system) and tested the associability of the cue in an aversive setting (defense system). To achieve this, we modified a serial prediction task (Wilson et al., 1992) that has been extensively used to investigate the neural substrates of associability changes in rats (e.g., Holland & Gallagher, 1993, 2006; Chiba et al., 1995; Bucci & MacLeod, 2007; Esber et al., 2015). In the original task, a light stimulus is initially followed by a tone that is partially reinforced with food (L➔T➔food, L➔T➔nothing). After developing an expectancy of the tone during light presentations, animals in the Surprise condition experience the unexpected omission of the tone on nonreinforced trials (L➔T➔food, L➔nothing), whereas control subjects continue to receive the initial training (L➔T➔food, L➔T➔nothing). The omission of the tone is intended to increase the associability of the light without fundamentally changing its predictive or incentive properties (which, if anything, should decrease during tone omission). This increase in associability is typically revealed in a subsequent test in which the light is paired with food (L➔food) and more rapid learning is observed in Surprise than control animals. Here, we tested the associability of the light by pairing it with foot shock in order to determine whether associability increases can be expressed across behavior systems.

Our results disconfirmed the hypothesis that associability enhancements reflect heightened plasticity within a particular behavior system. Rather, they are consistent with the view that such enhancements result from increased attentional processing of the cue (Mackintosh, 1975; Pearce & Hall, 1980; Pearce et al., 1982). Our procedure will provide neuroscientists with a tool to dissociate the neural bases of associability changes from those of other, behavior system- or valence-specific changes that a cue representation may undergo during learning. In the clinical setting, our findings suggest the possibility of administering associability-boosting treatments to bolster counterconditioning-based interventions.

## Method

### Subjects

Thirty-four experimentally naïve, male Wistar rats were used in the study, run in three cohorts that included animals from all three groups. They were obtained from the Animal Production and Experimentation Center at the University of Seville. Upon arrival, rats were acclimated to the colony room for two weeks with free access to food and water. The colony room was maintained on a 14:10 light/dark cycle schedule at a constant temperature of 21°C. Rats were housed individually in standard clear-plastic tubs (35×20×20 cm) with woodchip bedding. At the start of the experiment, they were 7–9 weeks old and weighed 230–280 g. One week prior to the beginning of the study, they were food deprived by progressively restricting their diet until they reached 90% of their original body weight and maintained at that weight thereafter. Once training began, they were fed a restricted amount immediately after the experimental sessions. They had free access to water in their home chambers at all times. All procedures and methods were carried out in accordance with the European Directive 2010/69/EU for the maintenance and use of laboratory animals and following Spanish regulations (R.D. 53/02013). The protocol was approved by the Ethics Committee for Animal Research of the University of Seville (Protocol Number: CEEA-US2015-27/4).

### Apparatus

Rats were trained in four identical, modular conditioning chambers (31.8 × 25.4 × 34.3 cm, Med Associates, Inc.) enclosed in a ventilated light- and sound-attenuating cubicle (63.5×41.9×49.4 cm, Med Associates, Inc.). An extractor fan was fitted on the right wall of the cubicle and produced a ∼60-dB background noise in the conditioning chamber. The side walls of the conditioning chambers were made of aluminum, while the front and back walls and the roof were made of transparent acrylic plastic. The floor consisted of 0.4 mm-diameter steel bars oriented perpendicular to the front wall and spaced 1.4 cm apart as measured from their centers. This floor grid was connected to a shock dispenser capable of delivering a foot shock unconditioned stimulus (US). Each conditioning chamber housed a 6-W white jewel lamp mounted 20 cm above the floor on the center panel of the left wall. Illumination of this lamp provided the visual stimulus used during behavioral training. A speaker was mounted 20 cm above the floor on the left panel of the left wall. This speaker was connected to a tone generator set to deliver a 1500-Hz, 80-dB tone which served as the auditory stimulus used during training. Each chamber also housed a recessed food cup located 2 cm above the floor on the center panel of the right wall. This food cup was equipped with an infrared sensor for detecting head entries and connected to a pellet dispenser capable of delivering 45-g sucrose pellets (DietTM; Mlab Rodent Tablet-45mg; St Andrews University). The chambers remained dark throughout the experimental session except during presentations of the light stimulus. In the same experimental room was a computer running Med PC IV software (Med Associates Inc., St. Albans, VT, USA) on Windows OS which controlled and automatically recorded all experimental events via a Fader Control Interface.

### Behavioral Procedure

Rats initially received a single session of magazine training in which a pellet was delivered in the food cup once every minute for a total of 30 minutes. They were then randomly assigned to three groups (Figure 1, table). In the first, Appetitive serial conditioning phase, the No-surprise and Surprise groups received Pavlovian magazine-approach training with a serial compound consisting of a 10-s light immediately followed by a 10-s tone. On half the trials, two pellets were delivered immediately after the termination of the tone (light➔tone➔food, light➔tone➔nothing). This training was intended to establish the light as a predictor of the tone. A Naïve group also received partial reinforcement training with the tone, but the latter was not preceded by the light (tone➔food, tone➔nothing). In each session, 10 trials were presented in pseudorandom order (reinforced and nonreinforced), with the constraint that no more than 2 reinforced trials could occur in succession. The mean intertrial interval (ITI) was 300 s. The mean number of magazine head-entries during the cues was taken as a measure of appetitive conditioning. Training continued for 10 sessions conducted over a period of 5 days, with two daily sessions run at 8 am and 3 pm.

**Figure 1.**
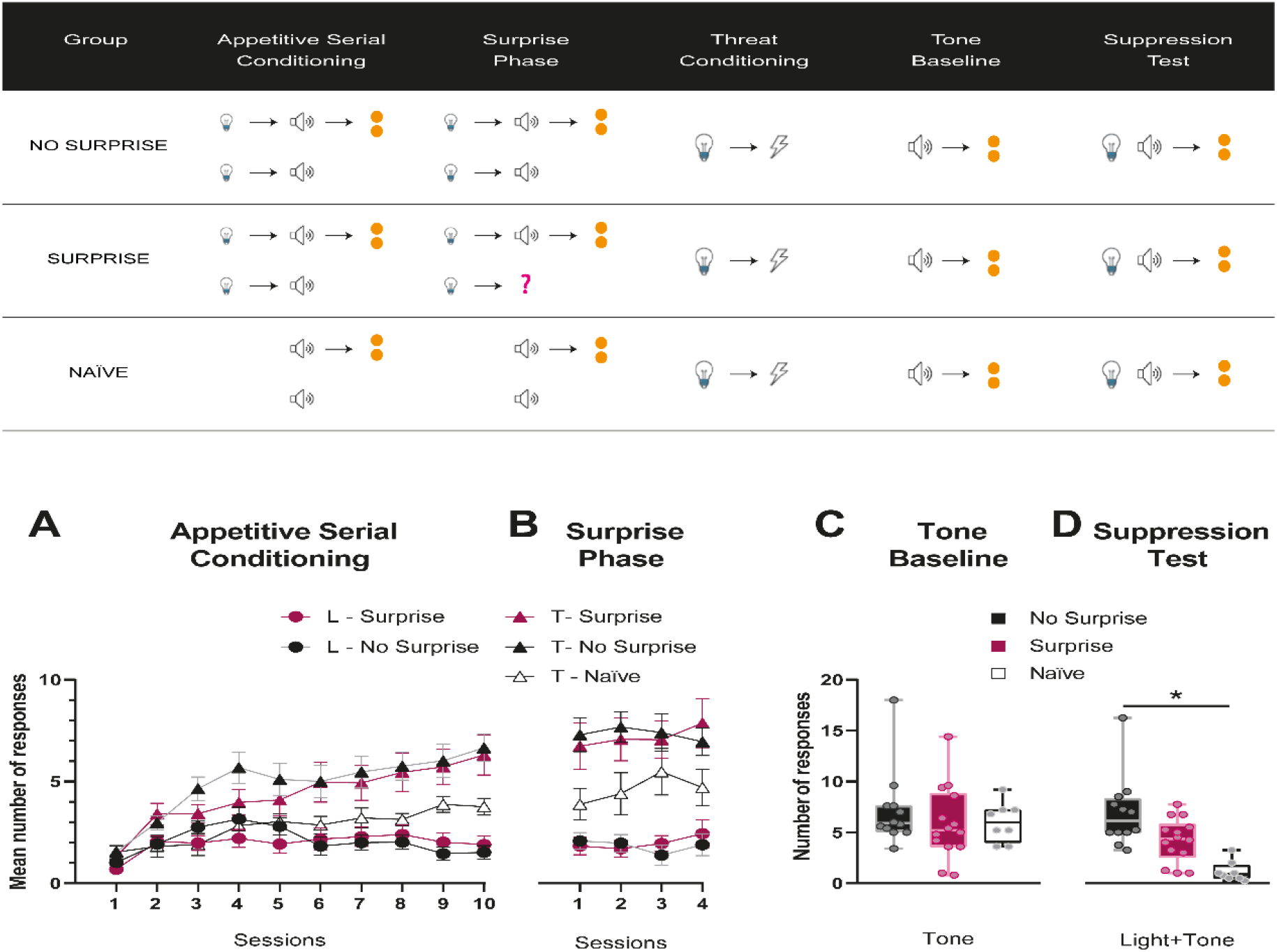
The table at the top shows the experimental design, a modified version of the Wilson et al. (1992) serial prediction task. Two groups of rats, Surprise and No Surprise, received appetitive serial conditioning in which a light was followed by a tone that signaled the delivery of food on a partial-reinforcement basis. As expected given their temporal arrangement, the tone evoked more conditioned responding than the light in both the Surprise and No-Surprise groups **(Panel A)**. A third, NaTve group also received partial reinforcement with the tone, but the latter was never preceded by the light. Following this phase, the No-surprise and Naive groups continued to receive identical training, but in the Surprise group the tone was unexpectedly omitted on nonreinforced trials—a treatment that has repeatedly been shown to enhance the associability of the light. Performance during this phase is shown in **Panel B**. All groups then received a threat conditioning session in which the light was paired with foot shock. Later that day, all rats were placed back in the conditioning boxes and given reinforced presentations with the tone alone to provide a measure of baseline responding **(Panel C)**. A subsequent Suppression test in which the light was presented simultaneously with the tone in extinction revealed greater suppression (i.e., more threat conditioning to the light) in the Surprise than No-surprise group, indicating that surprise-induced associability enhancements can cross behavior system boundaries. As expected, the greatest level of suppression was observed in Naïve rats, for whom the light was novel during threat conditioning **(Panel D)**.

In the next, Surprise phase, the No-surprise and Naïve groups continued to receive the same training for an additional 4 sessions (conducted over 2 days as in the first phase). In the Surprise group, however, the tone was omitted on nonreinforced trials (light➔tone➔food, light➔nothing) in order to boost the associability of the light (Wilson et al., 1992). All other procedural details remained the same in this phase.

On the next day, at 9 am, all groups received a threat conditioning session in which a single presentation of the light was followed by a 0.25-mA foot shock (L➔shock). This trial was preceded and followed by a 300 s period. No responses were recorded on this session. Later in the day, at 4 pm, rats were placed back in the conditioning chambers to receive an appetitive session consisting of 5 reinforced trials with the tone. The mean ITI was 300 s. The purpose of the latter session was to extinguish contextual threat conditioning and provide a baseline of magazine approach to the tone across the groups ahead of the final test.

On the following day, at 9 am, all groups received a suppression test consisting of 4 trials with the light and the tone presented simultaneously and reinforced with the delivery of two pellets (LT➔food). The purpose of this test was to measure threat conditioning to the light by assessing the extent to which it was capable of suppressing magazine approach during the tone relative to the tone baseline taken on the previous day. If the unexpected omission of the tone in the Surprise group enhances the light’s associability, and if associability changes can cross behavior systems, then greater suppression of responding to the tone should be observed in that group relative to the No-surprise group. The Naïve group provided a positive control for associability since those animals experienced the light as a novel stimulus on the threat conditioning session. Thus, we expected threat conditioning to be the strongest in Naïve rats.

### Statistical analysis

Analyses were conducted in R version 3.6.1. Generalized Linear Models (GLMs) were conducted using the *stats* package, Generalized Linear Mixed Models (GLMMs) were conducted using *lme4* package. To assess magazine approach performance in the Appetitive Serial Conditioning and Surprise phases, we collapsed (summed responses) across the last two sessions of each phase in order to probe asymptotic behavior. Before running any statistical inference, we selected to proceed with this contrast (the sum of the last two sessions) since the progression of responding across session was not of interest. We conducted all analyses with a single Generalized Linear Mixed Effects Model, adopting a Poisson as the conditional distribution of our outcome given the random effects and the covariates. We included a random intercept for each rat. We conducted post-hocs with a Bonferroni correction. Post-hocs analyses were conducted with the *glht* function in the *multicomp* package in R. All statistical tables are shown in the Supplementary Materials. Data as well as code to reproduce statistical analyses and tables are available at the github repository: https://github.com/gloewing/marquez-et-al-2021.

To assess magazine approach performance on the Tone baseline and Suppression test phases, we proceeded by collapsing across trials. We opted not to conduct a repeated measures analysis and no analyses of that kind were ever inspected. This was motivated by the fact that the progression of responding across trials within the test day was not of interest and thus the associated loss in statistical power from the increase in parameters we would need to estimate was not justifiable. The temporal nature of the data (i.e., the trial-specific structure) was a nuisance needed to probe the impact of the behavioral task, but did not provide any meaningful or interpretable information. Before conducting any statistical inference, all analyses were planned to avoid having to conduct any adjustment for multiple comparisons and to ensure analysis results were not selected to maximize statistical significance. Any comparisons of models were conducted without viewing p-values, confidence intervals or otherwise. Moreover, models were parameterized to provide the comparisons/contrasts of interest and thus no post-hocs were necessary. As such, Naive - No-surprise comparisons were not conducted. To assess whether there were differences in observed rates of head entries across the entire test day between the three groups, we conducted a GLM adjusting for baseline responding. Specifically, we used a negative binomial with a log link to account for potential overdispersion. During model building, we fit Poisson, negative binomial and quasi-Likelihood approach (quasi-Poisson) models and before examining p-values, compared models based upon the degree to which it accounted for overdispersion. To determine whether the model accounted for potential overdispersion we inspected fitted values vs. squared Pearson residual plots and conducted the appropriate likelihood ratio test using Pearson residuals. The only model that did not reach a statistically significant test for the presence of overdispersion was the negative-binomial model (Tone baseline: p= 0.255; Suppression test: p = 0.0996) and thus we based inference off this model for both phases.

## Results

Panel A of Figure 1 depicts the mean number of responses to the light (L) and tone (T) cues across the 10 sessions of the Appetitive serial conditioning phase. As expected, based on the serial arrangement of the cues, asymptotic responding to the tone (i.e., last two sessions) was significantly higher in the Surprise and No Surprise groups than that to the light. Surprise and No-surprise rats responded to the light at a rate that was, respectively, 67.4% (95 %CI: [64.1%, 70.4%], p < 0.0001) and 76.5% (95 %CI: [73.6%, 79.1%], p < 0.0001) less than they did to the tone, conditional on animal specific random intercepts. There were no statistically significant differences between Surprise and No-surprise animals in their rate of responding to the light (p=1) or the tone (p=1). Likewise, no significant differences in the rate of responding to the tone were detected between the Surprise and Naïve (z=1.563; *p*>0.709), or the No-surprise and Naïve groups, conditional on the random effects (z=-1.987; *p=*0.282). The numerically greater rate of responding to the tone in the Surprise and No-surprise groups relative to the Naïve group, however, is likely explained by the fact that the tone was signalled by the light in the former groups, allowing the animals to prepare for its arrival and respond at the magazine at cue onset.

In the Surprise phase, conditioned responding to the cues proceeded in similar fashion in all groups (Panel B, Figure 1). As in the previous stage, the Surprise and No-surprise groups responded to the light at a rate that was substantially less than that to the tone (71.4% in Surprise rats, 95 %CI: [68.7%,73.9%]), p <0.000, and 77.2% in No-surprise rats, 95 %CI: [74.6%,79.6%], p <0.0001). There was no significant difference between these groups in their responding to the light (z=1.192; *p*=1). Likewise, no significant differences were detected between these groups, or between either of them and the Naïve group, in their rate of responding to the tone (Surprise – Naïve, z= 1.797; *p*>0.434; No surprise – Naïve, z= -1.622; *p*=0.628).

Following the threat conditioning session with the light, reinforced presentations with the tone during the Tone-baseline phase (Panel C, Figure 1) produced no statistically significant differences between the Surprise and No-surprise groups (z= 0.992; *p*= 0.321), or between the Surprise and Naïve and groups (z= 0.076; *p*= 0.940). Crucially, in the subsequent Suppression test, greater suppression of magazine activity during the light/tone compound was observed in the Surprise than the No-surprise group (Panel D, Figure 1). Indeed, adjusting for baseline responding to the tone, the No-surprise group responded 41.8% (95%CI: [7.6%,86.9%]) more than the Surprise group (z= 5.270; *p*= <0.0001) across the entire test session. This difference suggests that the unexpected omission of the tone was effective in increasing the associability of the light, and that such an increase facilitated threat conditioning with that stimulus. Interestingly, however, the surprising omission of the tone did not fully restore the light’s associability to its original (novelty) levels, as suggested by the even greater suppression of magazine activity observed in the Naïve group in the test. Adjusting for baseline responding to the tone, the Naïve group responded 71.9% less (95%CI: [81.5%,57.1%]) than the Surprise group (z= -5.886; *p* <0.0001) across the test session.

## Discussion

Here, we employed a modified version of a serial prediction task (Wilson et al., 1992) to examine the scope of associability changes to predictive cues during learning. Specifically, we violated the expectancies generated by a light serially conditioned with food (feeding system) and tested the associability of this cue by pairing it with foot shock, an aversive exteroceptive stimulus engaging the defense system. To our knowledge, this is the first demonstration that surprise-induced associability enhancements can be expressed across behavior systems.

Our findings carry theoretical implications as to the nature of associability changes. From a behavior system’s approach (Timberlake, 1993; 1994), the results may be explained in terms of the close relationship between the feeding and defense systems. For many animals, foraging for food implies increasing their exposure to predators, making it essential to simultaneously attend to signals for food and threat. Crucially, preys and predators may share common predictive cues (e.g., a glimpse of a moving object, a rustle in the undergrowth), and thus it makes adaptive sense for associability increases to food cues to also benefit learning in connection with threat. It remains unclear, however, whether surprise-induced associability increments may be universally expressed across any arbitrary pair of behavior systems. Such is of course the prediction of attentional theories of associative learning (e.g., Pearce & Hall, 1980; Mackintosh, 1975). From this perspective, associability increments reflect augmented attentional processing of the cue rather than intensified associative plasticity within a particular behavior system. This assumption is supported by evidence that the same uncertainty conditions that foster new learning also promote stronger overt attentional responses (orienting) to the cue (Kaye & Pearce, 1984; Swan & Pearce, 1988; Beesley et al., 2015; Luque et al., 2016; Easdale et al., 2019). Notably, the fact that associability enhancements can be detected days after the induction of uncertainty suggests that the underlying attentional mechanism relies on enduring changes in the mnemonic representation of the cue rather than transient increases in arousal.

Critically, our assertion that associability enhancements can be expressed across behavior systems hinges on the assumption that the light—the target cue in the current study— was able to gain access to both the feeding and defense systems. One question raised by the low level of magazine approach evoked by the light in the Surprise and No-surprise groups during the first two phases of the study is whether this stimulus was capable of engaging the feeding system at all. That it did so is suggested by the greater level of responding to the tone in these groups relative to the Naïve group, which, as mentioned above, is readily explained if the light alerted the rats of the imminence of the tone and prepared them to respond at the magazine. Similarly, one could question whether the greater suppression of magazine activity observed at test in Surprise than No-surprise animals truly reflects a stronger activation of the defense system by the light or simply its greater proclivity to elicit competing orienting responses that interfere with magazine activity. While our study does not directly address this possibility, it should be noted that, by the same token, one would expect such competing orienting responses to equally disrupt rather than facilitate conditioning when the light’s associability is tested by pairing this cue with food. This is of course the opposite result to that observed in the standard serial prediction task on which the current study is based.

Due to various advantages, the standard serial prediction task has been extensively used to characterize the neural substrates of surprise-induced associability enhancements. One such advantage is that it permits decoupling the encoding of associability increases at the time of surprise induction from the expression of those increases at the time of learning. This advantage has permitted the discovery, for instance, that the central nucleus of the amygdala (CeA; Holland & Gallagher, 2006) and the substantia nigra pars compacta (SNc; Lee et al., 2008) are critical for the encoding, but not the expression of associability increases, although this has only been demonstrated in appetitive conditioning with food. On the other hand, the substantia innominata/nucleus basalis magnocellularis (SI/nBM; Holland & Gallagher, 2006), the secondary visual cortex (V2; Schiffino & Holland, 2016), and the dorsolateral striatum (DLS; Asem et al., 2015) are necessary for the expression, but not the encoding of associability increases. Such associability expression, however, has only been tested within the same behavior system (feeding), and thus it is unclear whether these regions would also be necessary in the current version of the task. Interestingly, the posterior parietal cortex (PPC), which has long been implicated in attention in humans and non-human primates (e.g., Mesulam, 1981; Posner & Petersen, 1990; Desimone & Duncan, 1995; Corbetta & Shulman, 2002) is so far the only region identified as being critical to the encoding, consolidation and expression of associability enhancements (Schiffino et al., 2014), suggesting it might constitute a locus for storing the cue-specific associability memory. Whether this mnemonic representation in PPC is fully detached from the motivational, emotional and behavior system-specific properties of the cue (i.e., whether it provides a neural substrate for the results observed here) remains to be established. Once again, the current procedure should help make this determination.

The present findings may also carry clinical significance. For instance, they suggest the possibility of expediting counterconditioning procedures by coupling them with associability-boosting manipulations (Keller et al., 2020). In behavioral therapy, counterconditioning refers to a collection of procedures that seek to modify maladaptive behaviors by associating their triggering events with an outcome of the opposite affective valence (Keller et al., 2020; Konorski & Szwejkowska, 1956). Recent studies show that counterconditioning therapies have greater efficacy (Engelhard et al., 2014; Kerkhof et al., 2011; Raes & De Raedt, 2012) and resistance to relapse (Kang et al., 2018) than exposure therapies based on extinction procedures. This relative advantage is thought to derive from the greater evaluative learning that takes place when the triggering stimulus is experienced not merely in the absence of its associated outcome (as in extinction), but in the presence of another of opposite affective sign. On the downside, however, counterconditioning typically requires more training than de novo acquisition or extinction (Scavio & Gormezano, 1980; Peck & Bouton, 1990; Bouton & Peck, 1992), and, to that extent, it could benefit from prior manipulations that enhance the associability of the stimuli and responses involved. The induction of surprise in a manner similar to that used here might provide one such manipulation. While our findings remain to be extended and replicated in humans, they point in a promising direction for future research.

## Acknowledgements

This research was supported by Spain’s Ministry of Science, Innovation and Universities grant PID2019-110739GB-I00 (to ED and JPV) and by the National Institute on Drug Abuse grant N5R00DA036561 (to GE). We declare no conflict of interest, financial or otherwise, related to the current research. We thank Felipe Cabrera for helpful comments on a previous version of the MS. All correspondence should be addressed to Guillem R. Esber.

## Author contributions

**Inmaculada Marquez:** Methodology, Investigation, Validation, Data curation, Visualization, Writing-Original draft preparation; **Gabriel Loewinger:** Formal analysis; **Juan Pedro Vargas**: Project Administration, Funding acquisition; **Juan Carlos López:** Conceptualization, Project Administration, Funding acquisition, Software, Visualization, Methodology, Supervision, Writing-Reviewing and Editing; **Estrella Díaz:** Conceptualization, Project Administration, Funding acquisition, Methodology, Supervision, Writing-Reviewing and Editing**; Guillem R. Esber:** Conceptualization, Writing-Original draft preparation

