## Supplemental Materials for "Surprise-induced enhancements in the associability of Pavlovian cues facilitate learning across behavior systems"

**This PDF file includes:**

Statistical tables

Phase 1

| | $exp(\hat{\beta})$ | 95% CI-Upper | 95% CI-Lower | z-value | p-value |
|---|---|---|---|---|---|
| Tone: Surprise vs. Naive | 1.403 | 0.918 | 2.145 | 1.563 | 0.709 |
| Tone: Surprise vs. No Surprise | 0.901 | 0.619 | 1.310 | -0.546 | 1.000 |
| Tone: Naive vs. No Surprise | 0.642 | 0.415 | 0.994 | -1.987 | 0.282 |
| Light: Surprise vs. No Surprise | 1.250 | 0.845 | 1.850 | 1.118 | 1.000 |
| Surprise: Light vs. Tone | 0.326 | 0.296 | 0.359 | -22.806 | $< 0.0001$ |
| No Surprise: Light vs. Tone | 0.235 | 0.209 | 0.264 | -24.633 | $< 0.0001$ |

Phase 2

| | $exp(\hat{\beta})$ | 95% CI-Upper | 95% CI-Lower | z-value | p-value |
|---|---|---|---|---|---|
| Tone: Surprise vs. Naive | 1.534 | 0.962 | 2.445 | 1.797 | 0.434 |
| Tone: Surprise vs. No Surprise | 1.030 | 0.683 | 1.555 | 0.143 | 1.000 |
| Tone: Naive vs. No Surprise | 0.672 | 0.416 | 1.086 | -1.622 | 0.628 |
| Light: Surprise vs. No Surprise | 1.296 | 0.846 | 1.984 | 1.192 | 1.000 |
| Surprise: Light vs. Tone | 0.286 | 0.261 | 0.313 | -26.982 | $< 0.0001$ |
| No Surprise: Light vs. Tone | 0.228 | 0.204 | 0.254 | -26.458 | $< 0.0001$ |

Tone baseline

| | $exp(\hat{\beta})$ | 95% CI-Upper | 95% CI-Lower | z-value | p-value |
|---|---|---|---|---|---|
| (Intercept) | 29.500 | 22.684 | 38.365 | 25.246 | $< 0.0001$ |
| Naïve (vs. Surprise) | 1.017 | 0.658 | 1.572 | 0.076 | 0.940 |
| No-Surprise (vs. Surprise) | 1.215 | 0.827 | 1.784 | 0.992 | 0.321 |

Test

| | $exp(\hat{\beta})$ | 95% CI-Upper | 95% CI-Lower | z-value | p-value |
|---|---|---|---|---|---|
| (Intercept) | 9.178 | 6.805 | 12.379 | 14.524 | $< 0.0001$ |
| Baseline | 1.100 | 1.062 | 1.140 | 5.270 | $< 0.0001$ |
| Naïve (vs. Surprise) | 0.281 | 0.185 | 0.429 | -5.886 | $< 0.0001$ |
| No-Surprise (vs. Surprise) | 1.418 | 1.076 | 1.869 | 2.479 | 0.013 |
